## Supplemental Data 1 for "Molecular characterization of nervous system organization in the hemichordate *Saccoglossus kowalevskii*"

**Supplimentary Table**

A)

| RNA *in situ* Gene | EST Access # |
|---|---|
| Dopamine transporter (DAT) | G142P60116RF4 |
| Glutamic Acid Decarboxylase (GAD) | G6027491317 |
| GPC | G178P60076RG2 |
| Histidine Decarboxylase (HDC) | G142P60145RF11 |
| Luqin (RWamide) | G142P60091RF7 |
| Prohormone Convertase 2 (PC2) | G826P59RB9 |
| Tryptophan hydroxylase 2 (TPH) | G826P5282RG11 |
| Tyrosine Hydroxlase (TH) | G613P6079RA4 |
| Vesicular GABA Transporter 1 (VGAT1) | G142P60044RE10 |
| VIG, IVG | G826P5251RB11 |
| GnRH | G142P60272RG7 |
| CalC | G826P5352RC9 |
| Vasotocin | G178P60059RH3 |

B)

| HCR Gene | Asseccion # | Sequence | Size | amplifyer | Probe size |
|---|---|---|---|---|---|
| TH | XM_006813504.1 | gCUAgACUggAggUCUAUAUUAUCU | 2890 | B1 | 20 |
| GAD | XM_002740628 | AUgUCUUUUgAUUUAggCUUCgAC | 4265 | B3 | 20 |
| VGLUT3 | XM_002739644 | AUgCCggUCAgCgUUUUUgAAAgUU | 1722 | B3 | 20 |
| DAT | NM_001168055 | ACACUggCUUAgUCgAAgUCACUUU | 3249 | B3 | 20 |
| Vasotocin | NM_001168017 | GTCCCTGCCTCTATATCTTTTAATAAT | 2250 | B3 | 33 |
| GnRH | XR_438542 | CAUAAAAAgAAggUgUCCAgAgCUC | 1500 | B2 | 20 |
| WFMRFamide | G826P521RB1 | CCCgACUACCAACUgACACAgAgCggg | 920 | B1 | 15 |
| NNFamide | XM_002739569.2 | UAAAUAUCCgCUAACgAUCCCAUAC | 1552 | B3 | 20 |
| CRH | XR_438216 | ggCAAAACAgAAAUgCAUUUgUgCA | 829 | B2 | 11 |
| orexin | XM_002734948.2 | CCTCGTAAATCCTCATCAAAGCTATCT | 1440 | B2 | 16 |
| CalC | XM_002735867.2 | CCTCGTAAATCCTCATCAAAACTGGA | 1800 | B2 | 20 |
| TRH | XM_002734602.2 | ACggAACACUgggAACAgAAgCUUgC | 1867 | B2 | 20 |
| achatin | XM_002732101.2 | CCTCGTAAATCCTCATCAAAGCGTAA | 2520 | B2 | 28 |
| CCK | XM_002738068.2 | CCTCGTAAATCCTCATCAAATTCTTTA | 1710 | B2 | 19 |

C)

| Primer Name | Sequence |
|---|---|
| flex_5prime | cgctttaattcaacctcaaaATGCTGTGCTGTATGAGAAGAACC |
| flex_3prime | gtccatgccgagagtgatccttACCCTCCGCCCAGGCCGGCGAACTT |
| flex_3prime-alt | gtccatgccgagagtgatccttCACCATCGACCCGAATTGCC |
| 5UTRsynapsin_flex | TTTTGAGGTTGAATTAAAGCGGGTGA |
| HLC5primer_new | GGATCACTCTCGGCATGGAC |
| HLC5Prime | ATGGTGAGCAAGGGCGAG |
| HLC3Prime | CGAACCACTTTGTACAAGAAAGCTT |
| 5utrTH | ctcgcccttgctcaccatTATACAGCCATAAAACTAATGTCAGG |
| 5th | aagctttcttgtacaaagtggttcgCCAAGTGAGCGAGTGAAACG |
| 5utrSynapsin | ctcgcccttgctcaccatTTTTGAGGTTGAATTAAAGCGGGTGA |
| 8Synapsin | aagctttcttgtacaaagtggttcgACAGTACATGCAGTTAAGGTGTGG |

D)

| Transgene name | Gene | Sequence insert | Vector Backbone |
|---|---|---|---|
| TH:eGFP | TH (XM_006813504.1) | CCAAGTGAGCGAGTGAAACGTATGC | HCL |
| Synapsin:eGFP | Synapsin (XM_006820227.1) | ACAGTACATGCAGTTAAGGTGTGGC | HCL |
| Synapsin:mGFP--T2A--(mouse)Synaptophysin-mRuby | Synaptophysin (NM_009305.2) | ATGCTGTGCTGTATGAGAAGAACCA | Synapsin:eGFP |

**TH:eGFP sequence insert:**

CCAAGTGAGCGAGTGAAACGTATGGCGATCACCTTTGGCCTTTCAGAAGGTTGCTATGCAGCTGCTTTA
GGAAATGCTTTGAAGATATTTGAGGTATGTATTATCCCCCAAAAATTATAGCGGGGGAGTCGTCACCTG
AGCATGCGTCAGATGGCCATAGCGTGTCATTGTTTACAAACAAGCGTCTACCCATGGGCCTTTGCGTTTT
TAGCGTGTGTTGCTATCGCCGCTTGGGGCTCAATTCCCGCGCGTGCGCAGGTGACGACCCTCCCGCTTTA
AACACATCGATACACAATCAAACATTTAACATCAGTTCAAATTTTATTGCCGGCAACATTTTGTTGAAAGC
CCTTTTATTTTGAATAACTTGGTCCAAGACGTTCTCCTATTCGTGTTTCGGTGATTTCATGACGCCAGAAG
ATATTGGACACTCATGAACAAATTCTACGGTCATTAATATCTGTTGCAATTGTGTATGACGTCACATTCCT
CAACGGGAATCGGGTCAAATATAGTTTTTTACGACTTTAACAAAGTGACTTTTTTAATTTTCTCAGCAGTT
AGAACCATTGCAATTTTTGATAAACAATTATGCCATTAGTGACTAAACGTGTAGTATGACGTGTTTAGCA
CCAACAAGTTTGCAGTGACGTCACGCGTAGTATGACGAGCCATGTCAAGAATGATACAAATCTGAGTAC
TTTGTGCCAAAATGTTTGCTCTGACGTCACGTGCACAGCGATGAGTCGTGCCACTGCTGCTGCAATTTTT
GCGGTAAACGACACACACTGTAACGTGTCACGCTTCAATCTTCCTGTGTATTGTAGACATTTCCCGAGGT
TCAAAGGTCATGTAACGTTTTCTGCTCTGCCTAAAATCGCATACGGCGGATATTTATTTTAACAAGAATTG
ATTTTCACTATGTGTTCTTAACCAAAACGTGCAAGTAACACAAATGCTAGCTAGAGAAAGAAAAACATAT
ACAGTTTCACTAATTATGTTTGCGTCAAACACCGCCCTTTCAATTCTCCGAATATCCGAGCTATTTATACA
GCGATGAATGTAGCTATTGTATTTTCATTTCCCGATTCGAATTGTGGAAAGACGTACATTGTTGCCTTGT
GCATTCACTGTAATTGACGAATCGTGGTCTACAATAATGATTCACCATGTTTAAGCGAAGTCCCCTGCGG
TCATTATGAGCAACGGGTAAACTAATTCAGATTTCCTTTTTAATTCTAGAATAATAACAATCCATTCTATA
CTTCACGTTTGAATAACATTGCATAGTGTTATCTTAATATAACATGGAAGCTCCAATATTGTCGAAAGTAC
CCAGGCTTACACAGCAACATCAACCACTACTCGTGATAGTTAAAGGGCGGTGGTTATCGCGCCGCGCAT
GTGCGACATTTGTGTTGATAAACATTGATTTTCCCATAATCCAATGCGCCTTACGTTACGGTGACAGATG
TATAATAGTAATTAATCAAAGATTTCTGCTTGCATATTCGCATAAGTTTTTGTTTTTTTGTTTTTGTTTTTTG
GATAACTAATCATTCATTTCACTCGATCATTGTTCATAATCCACTCGGGTTGAATTATTCTACAACTATTAC
TACTGATAAATATGCTGGTTTTTGCTACAAGTATAAACCACTTGGCACTTTTTTCGCCCATGTCTCTTGGC
GACCAAATTCTAATTGTTCGCTTTATGATATCACGAACTAATTTGTATAGCAGCCTACAATGGTCTAAATA
TTACTATTCTTTTGTATATTTTTCTTGGGAAATGAAGTTCAAATATTTGATCTTGTCTATTTGTACTCTAGT
GTAACAAACTGACTGGAGTTATATTTACTTTCGAACCCAACGTGTCACGTGGCACAGTCATCGTGAAGCG
CGTGTTCGAATTTCGTGAGCACGTGAACTCTTTACGTCACCTGGTGGTATATCACTTGAAAGCCGTGGTC
AAATTGGCCTACATTGTAATATGCAAATAACTTGTAAATAAGCCAATCAGCGCACATCCCTTCATACAGA
TAACCAATCAATCCACTTGAAAAAGCAGGCGATCCTTGCTCTAGTTTTAAACAAGCACTGATTGGTCATT
TGCTTTCATGACGTTTACTCATGATTACTCAAAACAATTCTCAAATATAGTCGAGAATGAGAATGCTTGA
AGCTTGAAGTAAAACATAACTAATACTCCGTAATCGCAACTTACAGTTACACTCAATGTAGATCATCGCC
AGGAGATGATGTCTTCACATCAGCGAACCGCCCCTTACAAGCATAGCTGTCCAACGCATAGTTATATTTA
AGTGGATACCTTTTAATAGGTTCAATATTTGTATAAACTGGCAAATGTTAAGAAAGATGGATGATATTAT
CAATCATATAATGAGTAATAGTGTAAATCACGGTTAATGTCCCCAGATTACCCACAAACCGAGTCAATAT
GGCCTGTGGTCATACAGAATAGACCATATGTAGTGCGGGTGTTTATGCCATAACATTTCGACAAACTAG
GCTCTAAATATCATACGGCATTTTACAGGTTAAATATGTTAAAAGGGCTGTCAGAAAATCCTACAGAAAA
CAAAATCTAGATACAGATAAAATAAAAGTCCAGCACGTGTTCGTATTCCGTGTAAATCAAAGATGCATTT
AGCACATTTAAACATGTCGAAGACGATTATTCGCAAAGACCATCACATTTAAATTAATTGTGGATAGCAC
ACTTAGAACACAGCGGCAAGTTCATCACGTAGTTGTTTGCCTGAAAATTATGCTATCCTGTCTCCATACTT
TCTCATTGCATGTATATACGCAACTGTTATTCCTTTCAACGGAAGTACATTTGTTACGGCACCAAACTGCT
TCACAGACCTGATATGTGGTGTCGGTACTTTGAAAATGAATTATATCTTTATTGAGCGAATTATCCTTGGT
TATTAAGGATTCTCCCATTTCAAAATAGTGGATTGACTGGAAGATGTTGCAGTGAAGAAAAAACAAATA

ACATGATTTTCAATTTCCATGACAACAACATAGCGTCGAATGTGAATAATAAAACTGACGTCAGCCATGG
GTGAATAAATATTTTCTAAATTCCACCATCCCCACTACCCACCCTGTAATCTCCTACAACGGAACCCCGCG
TATAGACAGGCTCATGACGTGCAATGTGGTTATAATGTAAATCTAAGAGAAACTGGAAAATCACCCTGG
TGTATAATAATAGGTCTTCAGGTGGAATTATCGGCTTATATGCGAATTTCCGATTACATCATTCATGGTAT
CCGCACGTGCCAGAACTACGTTTGCGCGGATTTGACAGGCATCTACAATGAAGAAATACTGTGAATCGA
GCCCAGTTCTGTTCCATTTCAAAAATCCCTACATAGAAGTTATGTGTTGGATTCCAAAATCGTATGTTTGC
CGGTTGTTTCCATGGCAGCGTCAATCAAAAAAGTCCCGCACGTGGGTCAATGTATGTCTGTGCTATTTAG
TTGGCTGTTAAGTAAAACAAACTGAGGAGAACACCGTGTGTATAACTGAAACCAACAAATATTGGCTTTT
TACGGAACTAGGGGGGTTCATATGATAGACGATATGACCATCTGTCTAATGTCAATCTAATACTGCATCT
GAATCAATGTTTACAAGACATTCATTCATGATAGTGTAAAGGATACATAGAGTAAGAGACGGGAGTACC
TAGTACATGGTATCAGTGCTACGTCGCAAGCGAGGTCAAAACTGGCGTATACCTTGAACAACCTTATATA
TTGGCCCTACATAGACGAGGTCAAGATGAAAAGTAATCCACAGTCATTTTTTTTCACGATTCCACGCAAG
ACTGGTTTTGTTTCGGTGTCACTTCCTGATTTACGTGGAGTTTCTCGTGATAGTGCATACGAGTGATAACT
TAAGTGTAGAAAATCGGGGATAATTACAGCTCAATGCAATCAGAAAACTGAATAATAGGCCACTTTTAA
TTCCATACATATAACCTATATTTAATTCACATTTATTCGAATGTATTATGATAAAATAAAGTTTATACTTTC
ATTTAAATCACTGTATTATAAGTTTACCATGATATGTTGCTAATTAATGGGTTGGGATGAATGCTTAGTAT
CTAGCGTTTACACCAGCTCAGTGAAAAGTCGGCGATTCACGGGCTCTCGCTTGGTCATGCGCGAGATAT
GCGCCTATAGCAACAAAGATGGCGACTTCGAGTATTTCGCGGATCAGCGGATACAAAAAAGATACGCTA
CCAATGTAATTCTTAGAATGATTATGACGTATATCAAAGAATATTTCAAAGTCCAAGTCGATATTCAGTG
CACATGACACGCAGATCAAAACTCGCGTTCGTGCTTTTCACCCTGGGAGCGCACTGTATTATGGGAATCT
CAACGTACTGTATATCTCGCACATGCGTGGCACGAAGACTACCGCCTTTTAGTAAACAGTTACAGCGTCA
CGACAGTCGCCGTAAAGCGCGTTCGAATTTCTTGTAATATAAATGTGCTGCAACAACTTAGGCTTTTAAG
TAGGAAGTCCATATTTGTAGTGTTAGGCAGTTGTAGAACTTCATTTCCTTGAATCAATCATACAAAACATT
GTTTGATTAGCGCAGCCTTGACGCCAATTTGTGGCATACCGTGCTCCACGAGGCTTTTAGTGTCAATTGT
TTCCTTAATATTTTTTGCCTTTGATTCTTACCTTGAGGGTTTCTTCAGATTGGAACGTGCTGAAAGACAGC
TACTTTATTATATACTCTTTATTTATGAACGTTTAGTGACGACCTAGAAAAGAGCTGTAAGGGTTTTTGAG
AAATGTGATCATAGAGCATCACGTGACGGAAGTGGAAATATGCCATTTGTAGCTCATTAATGACAAGGA
TATTTCTAATTCGGCTCACAATTTTGTTGATTATTACGCAGACTAATATAGGGCGAAAGCAACATGTTGA
ACAGAGTGTGTCCATTCTTTTCGACTGACACCTGACATTAGTTTTATGGCTGTATA

**Synapsin:eGFP sequence insert:**
ACAGTACATGCAGTTAAGGTGTGGCCTTATGCTAGCAGTCATTTCTTAGACATTATTCACGATATGGGCT
TGACATTTTAACTACATTTGTATCAAGATAATGAAGAAAAACCTTTTTAATTTTTTTTTATATTGTCATTAA
AGCTGAAACAATAACATTTAAGAACAGCCATTGGGAGAATGTATAATAATTTTGAATTGTTGGTATGTAT
ATTAGGCTTGGTGTCTGTTCTGTTTTAAAAAATATTTAATATTAAAAATATGAGAAGACTCATACGACTG
ATAAATATAATGGTAGTAGGAAATCTTAATCTTACCAATTTCTATTATAGATTACACTTAGAATATTAATG
TTTTTAACTAGATTATAATAATGAATAATGTTATTATAATTAGTCATAATGTATATGAATATAAACTTAACT
CCTGAGAGATTTTCATAATAAATAATCAATTGATCGATTTGGAGTTTAATTTGTTTTCAATTATCACAAAT
GTTTTAAAGGAAACTTAATATTACTTTGTCCTTATTTCAAAAAAAGTGAAAGTTACTGAAATGTTCAGTTG
TGATGTATGGGCGCTATTGAATCAGTCATGCACAACATCTGTCACATGTAATTACTCTCTCATTAGTACAA
GCACACCTGCTGGCTTTGAGGTCATTTCCCATCATGCAACAGTTCACTATATGATCTTAAGGTGTGTATAT
CTTGCTATAAAAAGAACAAAAGAGTGATTTTTAAGTTGTGCTATACATTACCAAACCACACCGAATATAA
ACTTTCGTTTATAAAGTAATAACCAATTACGATGTGTTGATGATAAATTACTGCTACGAGACGAGTCTGT
CCCATGGGAAACAGAATATGTACAAGAAACATGTAGCACTGACCTAGAGTTCACTATTGTATGGATTAG

AATAAACTTTAACAATGCTTGGTTCATTATTAGCAGTTGCAGGTGAGTTAAGATGGATATAATATCTTTA
CTTGAAAGAAGGGTTTATTAATTAAGGCCTTTAATGCTGTTCCAAAATGTTGTCACCATAGGTCTGCAAC
TTGTATATAGTATATGTTCAAAGTTGAGAGTTTATTAGCAAGCAAATTGAACGTAAGGTTATTCAGTTCA
CATGTATTAATCTGAAAGGTGCATTATTTACTTGTACATTGACTGTAAAAATGTGTCTAGAATTTGTTAGT
TATTTTCAGAAATTCACAGAAAATTAGTGTTTTATTAGTGTTAGTTTTTCACATTTTGTCACTAATCCTTCA
GCACAAGATGTTCTACTTGTAATGCCTTTGGGAAATCTCCAAACCGACCGTTCTAAATGTGAGCAATGAA
GTTCTAAATGATACACATGAAAAACCTTTAAATCTATGTTTTAGACTTAGCTGTATAGAATCTGATCTCCA
CATGCCCATTTTTGGGGCAAAATTTAAGTGCCTCAGAAATTGGTCAAAGCTCCAGAGGGAAAGAATCTT
GTTTCATATAAGGGGATTTTGTCAATTTCATGTGTTTCTTCATTGTGTGCTACTGTGTCACAGACTAATAA
TAAGACATATTAATACCTACTATCGTTAATCTTTGAATAAAAAAAAATGATCAAATGAAAGTGATTTTGA
AAGTTGACAATGTCAGGGAAAGGGTCACTCTTATTGATAATTAATTTTGCAAAAGGAAGCAATTGTGAA
TTGGAAAATGGCAAAATGATGTAGCAGTGAAATTTAACGATGTACAGTATAATTAGAATTTTTTTGATAT
TCTTTATTTATTGCATTGATGTCAAAATACCATTGAGGTGGTGAACAAGGCCTTTGAGGTCTCAATACAA
AATTTGATTTTGTGCATTTTCAGTAAACTGGGGAATTGTATTTGCATTAAAACATTGAGAAAATGAATTAT
TCATGTTAAGTTTCCTGGCATGTTATTTTTATTTCAGAATAAATAATGCAACAGACATATCAAGCAAATAT
GCTTGATATAACTGAGGCTTAAAATCCAATATGCATTTTAAATATTAAATTTGCAATTCATTCACAAGCAT
AGGAAAATATACATGATAATTAATGTGTAAGATTGTATTATAGGATTACAAATTGAAGACTTGATTTTTA
GCATTGTTTTTGCACTCTTGCTTCATTTTTGAAAATAACTCCCCTGTTATTGGGCCCCATCAAATAATTATC
TCAAAATGAATTTCATATATTACGTTAATTCACACCACATACAATGAAATATATTAAGTGCCGGTACACTG
CCAGCGGTTCTCAACTTTGTTTTGTTCAACTATATACTATATAGTATATATTGCAATTTGATTGTAAGGTTT
CCTGCAGAATCCAACATGATGAATCAATTAATATACACTTCACTTCAATGCAGGTACTGTACATTAAATTG
ACAGTACATGCTTTCCCGTTGACTCAAATGTGATGCATGATTTAGAATAGGAACATCCAGGGTATTTTGA
GGTCTAATGAAGTGACCTTTATTTGAGCTGAAATTGCTAAAAAATGTATTTTGGCTGTTTTGTATGGCATT
TATTTTAACTGAATTTTGATGAGTTGAATGTAGAACACAGGGAAAAAATGTGTTCACTAATTCATCCCTTC
AGCGAAAATAGCAAAAATAAATTCCAATTATTAATAATAAAGTATTTTACATTCTGGAGAGCTACCTAGA
AACATAACATAATACTGGAAGTGCATTAAATATTCACTGGGTTTTAGTTATTAAGTTCATTGTTATTTTTG
CTGATGAGTAAAACCCGCTAAAAAATTCACATGGACTTGTAAAATTACAATAGAACTCATTATTTTGCTG
GTAATTCTGCGAATTTAAAATTCAGTCAATATTCAAAATGTACAAATAGTGAAATTAAATGCCAGCAAAA
ATATATGGTTTTACGAGACTAAGTAACTTAATTGAAATATGTACTTTTGTACCCTTGTGCAACTATCATGT
TTTATATTATTGCAACACTTATATCATTTAAACTTATAAGTAAATTGGAAAGAATATGAATATATGAGTAG
AGGTCGATGCGATTGTCTACTGTCTGACTGCATACTTCAGGGTATCCTTGTGATAACCTTATTTTCACTGA
CTTACGAAATTAAATCAGCTTTCAGCATATTCACACTTTGTGATAGAAACAAGAAGGCTAGTGTAGCTGT
CTCAATCAGTGTATTCTCCACCCACACAGATTTCTCTTAATTTATTGAATAAATAGGAATAGTGAGTGGTG
TTAACTAAATACACTGATGAGATTTCTTCCCCCAATGGGATTATATACAGTTGCACCAGTGAATGCAAAA
CTGTTTCATTTAAAAAAAAAAATTGCTAGATTACATATATTTCATATATTAGCTGACAAGTATTTTTTACACT
ACTGTGAAATACTTTATTTTAGCTGGAATTAATTTTCGCTGAATAGACTTTTTGGGTGTTTTGTATGGAAT
TTATTTTCACTGATTTTGATGAGTTCAGTGTAAAATATAAGAAAAAAGGTCTATTCACTGAATTTTATTTT
CACTGAATCCTTCACCAAAAATAGTGAAAATCAATTCTAAGCGAAAATTAAGTATTCTATAGTACCGGTA
TATTGTATTATTTTTCAAACCACTTACATACAATTGTTATCCCTTGAGTTATCAGAGGCCTGCTGTTACCAG
TAAGAGCAGCATTTGTGGTCATGTGACTGTGATTATGATATCCATCTACTGTGTCCTGGACAATTTTAGG
TTGGCTGTCATTGAATGAATCTAATTATGCATAAAATCAATCAATAGTTTCACGACTTTGCTTATCAATTG
AGCGCTAAATTCTTCCATTTAACGGGTCTTAGGTCTTTATGTATAATTGTGTGTTCATAAAAGAATGATGC
TAGGCTTTATGTGTGACAATTATTGCTCACATAAAGAAAAACTACCATTATCATAATTTTACTAATACTGA
AGCCCAAGCCCTGGTGTATATTGTTCAATGTATTCCTAGATAAACTACATGTATATTTGTTGACTGTTATT

AAGCGTACCAAAATATGTTTTTATGCTTACCTCATCAGATTTCAATGAAATTTTGTGCACTGGGTGATATG
GGGATAACTGAGTGACCAAAATAGTTCTTTCTTTTAATCTATGCATATTAATTATCTAATTTGTCTAATAAT
ATTACCTCAAATTTATATTCAATGGATGTTCTCATTAGACTATAAAGGAATTCATTAGAGACTGGGATTTA
TGTGATGTGTGATTTTTAACATCAAATCTTTGCAAAGTAATTAGTAAACACTGTTTTTTTATAGAAATATA
TTTATTATGAACTACTTGTAGCTAATATGCTAATTTAAGTTATCAAACTAATAAAAGTTAAATATTTTATTT
TACCTTCAACAAGCTAATTTTCACCTTTGTAATATCTAGTAAATGAATTAATTGTATTTAGAATTCAATTTA
ACCCTATTAGAATAGTAGTATATATTCGTCCATTACAAAATAGTCGAGCCAATATAGTTTCAAAATGTTAT
AGCAGATTGATGATTCTTGATAATATTGAGGTCAGTACATGGTAAAATATTTGCCATCTCTAGATTTCTCC
ATTTTGTAAAATCTTGATGACAAAAATTGGATTTATTGATAATCATCTAATTTGATCTTTATCTTGTGTCTT
GTTGTAATGAGTTTGTTATTACCGGCTCACAATTGATGTGTATGTGTCATCTCTGAAAGTCACAAATACTG
ATACTTTTTGGCATATTGATGCTCTATAAAAATACTTTCACCTGGAACTATTTTCCTATATTTTACATCAAA
AATCAGTGAAAATAAACTTCATACAAAACTGTCAAAGTGTTATTTTCAATGAAAAATAAAATATTTTACAC
TACTTCAATTTTTGCCATGTACTGTTTTTGAATTTTTATGATTACTGCTTTGGGGAAAATAATTGAAAAATT
CTCATTTGACCTTGGTGGACTTTGTATTTTGGATTCCATCCCTGCAAATCATAATGGGATATCCATATGCA
CATCAATTTGTTTGGTTGACGTGTGGCTGACTGTCAGCCATTTTGTCCCAAATTACATCTAAACTGTTTCT
CGGGCATACCTGGACTGATTTCATTGTGCCATTGACATTGGTATTTTTTATGTAAGTCAATTGGTTCACAA
TTTGATGCATTATTTTTGTTATAAGTGATACACTTTCACAGATTTGTTTAATTTTGGTACAAACACAAGTCT
TAATTGGTTGCATGAATTGGTTTAGTTTTTTAAATAGTGAAATTACATACATACTGTGCCAAGTAAGTTTG
TCTTTCTCCCATTCTGCCCAATCACTGTCAAAATTCAGTAGCACGTACACCTCCCTAAACTGCCAGTCAGT
TGATAAATATTAATTAGTAAGTGCATTTTTGTACCCTATTTGCATTTTAATTGTGAATCAATCCTTTTATGG
GCATGCCGGTAATGACTTCCCTGTTTAGGTCATAAGAGATTATTTTTAAAAAAGTACCAACATAGACTCT
GTCATTTTCAATGACTTGATAGTGAATTAATTGTACTTTCAGAGAGTACCAGAATGTGTGATCTTTCGGA
GATCATGCATGAGTACTATTGCCCACTCGGTCTGTATGGCATTGCGTTTCTTCCCAATAATCAAACACAG
GTAGAGATGTCACTTCATTCTCAATTTAAAGGTTAACATGGTAGTGCTTAATTGGCATCACATCTCATGG
GATTTAATTATTATCTATGAGTGTTCAATAGTTACAGTGTATGTGAACTGTGTGAAAATGTGACTGATCA
GTTTATTCAATGTCTATAATATAAATGGTGCAACAGTAATGTAATACTGATTTATCACTGAGTAAAGGAT
TCAAATCACAAAAAAATTTGTGTCTCTTACTTTTAAATCACAAATTTTGATTAGCTCCACTGTCGACTTTGTC
GAGCGGAGCTTATCATATGAGTGATTGTCCGTCGTCATCTGTGGCAATGTTTTACTTTTTCGACTTGAAA
ACTACTAGCCTGAATGCTTTGATACTTGGTGTATATGTACCTTGGGTAGACCTCTCTTAGGTTTGTTCATT
TCATGTTGATATCTTCAATTCTCTAATTTTGGTGTTTTGTTAAAAACCTCAAGTCTAATTGCTTTGATATTA
TATATATAAGTACTACATGTGGACTCTCTTTCAGATTTGTTAATTTCGTGGTGATATTTTCAATTAGCTAAT
TTTGATGAGTATTTTTGTCATTGTTGGTGAAAAGTGATATTTTTGACTTCTTGTTGAAAACAGCAAGTCCC
ATTGCTTTGATACTATACATATTGTTTATTTTCTAATGACATCTTCTGTTCTTTAATTGAAATGTGCCTTGCT
AACAGCAGAGCTATCTCGGCCGACCTGCTGCTTGTTTTGAATATACATTCTGTAAATATCAGATACTGTTT
TCATATTTGTTTATTTAATATTTATAATAAATGCAAATTAATTATCCTCATTGAGACATGTTCTTGAAAATT
ATTATTATAGTTTTTTGGTATTCAATGTTGAGATACAACCTTGGCATTTACATTAAATAATTGATTAAATA
GCAAAGAGGTCAGTAAGTTATTATGAATGGAACATTTCATATTAACTCCATTAAAGGAAATATTGATTGC
TATTTTCAGCAAGTCAGAATATATTATCATAAAGGTATTGAAAATTTAAATATTAAAATGCACTAATAACG
TTCATTATTCCATATTCTATTGCCAAAATTTCTAATCAGAATTGGATGGAACAGTCAAACAGAAGTAATAT
AGACAGCAGTAAATGCACTCTGCTTAACAATTTGTTTTGCCTAAGGTCTGCAAAACACATATTTACTGTTG
AAGCTCATTATAATACGTCTGATTTATTTTCTGATATGATCCAGGTTTTTGAATAGTTCAATTTATGAAAG
ATAATCACTTCATTATTTCAGTGCTTTATTGCATCTCAATATGAATAAATTGAATTTTTATTTCTAAGGGAT
TGGAGAAATACTTCACTCCAGATGCTACTAGATTATTCTTACTTCTGAATCACTTTGGAGACTGTCAGTAT
GAATTTGAAATTCTTGTGATATTTATATATCATAATAGTCATTAATATTTCAATATACAGTATACTTTATAT

TCAGATATATGAAGGATATATTTTTTATTACTGATAATAATAGAATGCCATGTCCTCGACTTGAGGTTTGT
GGGATATGGAAATTATTTTCATGAGTGATTTGGATATCCTGCAGCAATATTTTCGCAAACAGAGTGAGTG
AAAATACAGCATGTATGCACCACATTACATATCTAATAGTATGACAGATAGTATATTTGTTAGTATTTTCA
CAAGTTATGCTGTAAGTGTAAAAAGTGAAAATAAAGCAATGGATTCACTGCAACACTTTCAAAATATAAT
TTTTTAGTGAGTATCTTTGAGTATTTTTAAATTATTAATGTTTGAGAAGATTACAGTGTATGTGAGGCGTG
ATAAATTTGGAATGGTACAAATGCTATATTCTATGCTAAAGTTTATTTTATCTAGTTGTCTGAATTGTGAC
ACCTTTGATAATCACTTCAAGTAAATCATTCCAGAAATACCGTAAAATAATATATTTTGTCTTTTTCACATA
CTCACTCACTTACATTTCTTTTTTTTTTCCAGACAAATGAAGTGTGTATGCTAGAAGATTGGATTCTTATCT
GAGTGATTCTAGGAATATTAATACCTAATGGAAACAATAGATTTTGTGTGATTCTACAGTATATAGAAGA
TTCAACAATAATTATTCAGCCAATCATCAGGGACCTAATTTACGCTTGGATGTACTGGTTTACAGATAAA
ACTGAAGTTGCGATACGATATAATCCTATATTTTTTTCATAGTAAGGTGATATTATTTTATAGTTGCGAAT
AAATATTTGAACTCTATTGCACCTGCTGTCGAGAGAGCTGGTTTTCTTATTTTTATGTGGCTTTGGTACAG
TTGATTTATACTCAGTAACCACAAGAACTTCAGCAATAATTTGATCCGTAGACAGTTTGTCTGCCGAGTG
TAGTCTTATTCCGTACTTGCTATTACATATTGGCCAGCTGTCAAAGAGGATATTACAATTGCGGGTCTACC
TCGTATAGTGAAATTCAATTCAACGCTGGCTTCTTATTGAACTAAAATCACCCGCTTTAATTCAACCTCAA
AA

**(mouse)Synaptophysin sequence insert:**
ATGCTGTGCTGTATGAGAAGAACCAAACAGGTTGAAAAGAATGATGAGGACCAAAAGATCATGGTGAG
CAAGGGCGAGGAGCTGTTCACCGGGGTGGTGCCCATCCTGGTCGAGCTGGACGGCGACGTAAACGGCC
ACAAGTTCAGCGTGTCCGGCGAGGGCGAGGGCGATGCCACCTACGGCAAGCTGACCCTGAAGTTCATC
TGCACCACCGGCAAGCTGCCCGTGCCCTGGCCCACCCTCGTGACCACCCTGACCTACGGCGTGCAGTGC
TTCAGCCGCTACCCCGACCACATGAAGCAGCACGACTTCTTCAAGTCCGCCATGCCCGAAGGCTACGTCC
AGGAGCGCACCATCTTCTTCAAGGACGACGGCAACTACAAGACCCGCGCCGAGGTGAAGTTCGAGGGC
GACACCCTGGTGAACCGCATCGAGCTGAAGGGCATCGACTTCAAGGAGGACGGCAACATCCTGGGGCA
CAAGCTGGAGTACAACTACAACAGCCACAACGTCTATATCATGGCCGACAAGCAGAAGAACGGCATCAA
GGTGAACTTCAAGATCCGCCACAACATCGAGGACGGCAGCGTGCAGCTCGCCGACCACTACCAGCAGA
ACACCCCCATCGGCGACGGCCCCGTGCTGCTGCCCGACAACCACTACCTGAGCACCCAGTCCGCCCTGA
GCAAAGACCCCAACGAGAAGCGCGATCACATGGTCCTGCTGGAGTTCGTGACCGCCGCCGGGATCACT
CTCGGCATGGACGAGCTGTACAAGGAGGGCAGGGGCAGCCTGCTGACCTGCGGCGACGTGGAGGAGA
ACCCCGGCCCCATGGACGTGGTGAATCAGCTGGTGGCTGGGGGTCAGTTCCGGGTGGTCAAGGAGCCC
CTTGGCTTCGTGAAGGTGCTGCAGTGGGTCTTTGCCATCTTCGCCTTTGCTACGTGCGGCAGCTACACCG
GAGAGCTTCGGCTGAGCGTGGAGTGTGCCAACAAGACGGAGAGTGCCCTCAACATCGAAGTCGAATTT
GAGTACCCATTCAGGCTGCACCAAGTGTACTTTGATGCACCCTCCTGCGTTAAAGGGGGCACTACCAAG
ATCTTCCTAGTTGGTGACTACTCCTCCTCGGCTGAATTCTTTGTCACCGTGGCTGTGTTTGCCTTCCTCTAC
TCCATGGGGGCCCTGGCCACCTACATCTTCCTGCAGAACAAGTACCGAGAGAACAACAAAGGGCCAATG
ATGGACTTCCTGGCCACAGCAGTGTTCGCTTTCATGTGGCTAGTTAGCTCATCCGCCTGGGCCAAAGGCC
TGTCCGATGTGAAGATGGCCACTGACCCAGAGAACATTATCAAGGAGATGCCTATGTGCCGCCAGACAG
GAAACACATGCAAGGAACTGAGGGACCCTGTGACTTCAGGACTCAACACCTCGGTGGTGTTTGGCTTCC
TGAACCTGGTGCTCTGGGTTGGCAACCTATGGTTCGTGTTCAAGGAGACAGGCTGGGCCGCCCCATTCA
TGCGCGCACCTCCAGGCGCCCCAGAAAAGCAACCAGCTCCTGGCGATGCCTACGGCGATGCGGGCTAT
GGGCAGGGCCCCGGAGGCTATGGGCCCCAGGACTCCTACGGGCCTCAGGGTGGTTATCAACCCGATTA
CGGGCAGCCAGCCAGCGGCGGTGGCGGTGGCTACGGGCCTCAGGGCGACTATGGGCAGCAAGGCTAC
GGCCAACAGGGTGCGCCCACCTCCTTCTCCAATCAGATGTCGGATCCACCGGTCGCCACCCGTACGATG

AACAGCCTGATCAAAGAAAACATGCGGATGAAGGTGGTGCTGGAAGGCAGCGTGAACGGCCACCAGTT
CAAGTGCACCGGCGAGGGCGAGGGCAACCCCTACATGGGCACCCAGACCATGCGGATCAAAGTGATCG
AGGGCGGACCTCTGCCCTTCGCCTTCGACATCCTGGCCACATCCTTCATGTACGGCAGCCGGACCTTCAT
CAAGTACCCCAAGGGCATCCCCGATTTCTTCAAGCAGAGCTTCCCCGAGGGCTTCACCTGGGAGAGAGT
GACCAGATACGAGGACGGCGGCGTGATCACCGTGATGCAGGACACCAGCCTGGAAGATGGCTGCCTG
GTGTACCATGCCCAGGTCAGGGGCGTGAATTTTCCCAGCAACGGCGCCGTGATGCAGAAGAAAACCAA
GGGCTGGGAGCCCAACACCGAGATGATGTACCCCGCTGACGGCGGACTGAGAGGCTACACCCACATGG
CCCTGAAGGTGGACGGCGGAGGGCACCTGAGCTGCAGCTTCGTGACCACCTACCGATCCAAGAAAACC
GTGGGCAACATCAAGATGCCCGGCATCCACGCCGTGGACCACCGGCTGGAAAGGCTGGAAGAGTCCGA
CAACGAGATGTTCGTGGTGCAGCGGGAGCACGCCGTGGCCAAGTTCGCCGGCCTGGGCGGAGGGTAA
GTCGACATAACTTCGTATAGCATACATTATACGAAGTTATGTGTCGATGGTGATGCTTGGCAATTCGGGT
CGATGGTG

Supplementary Movie 1:

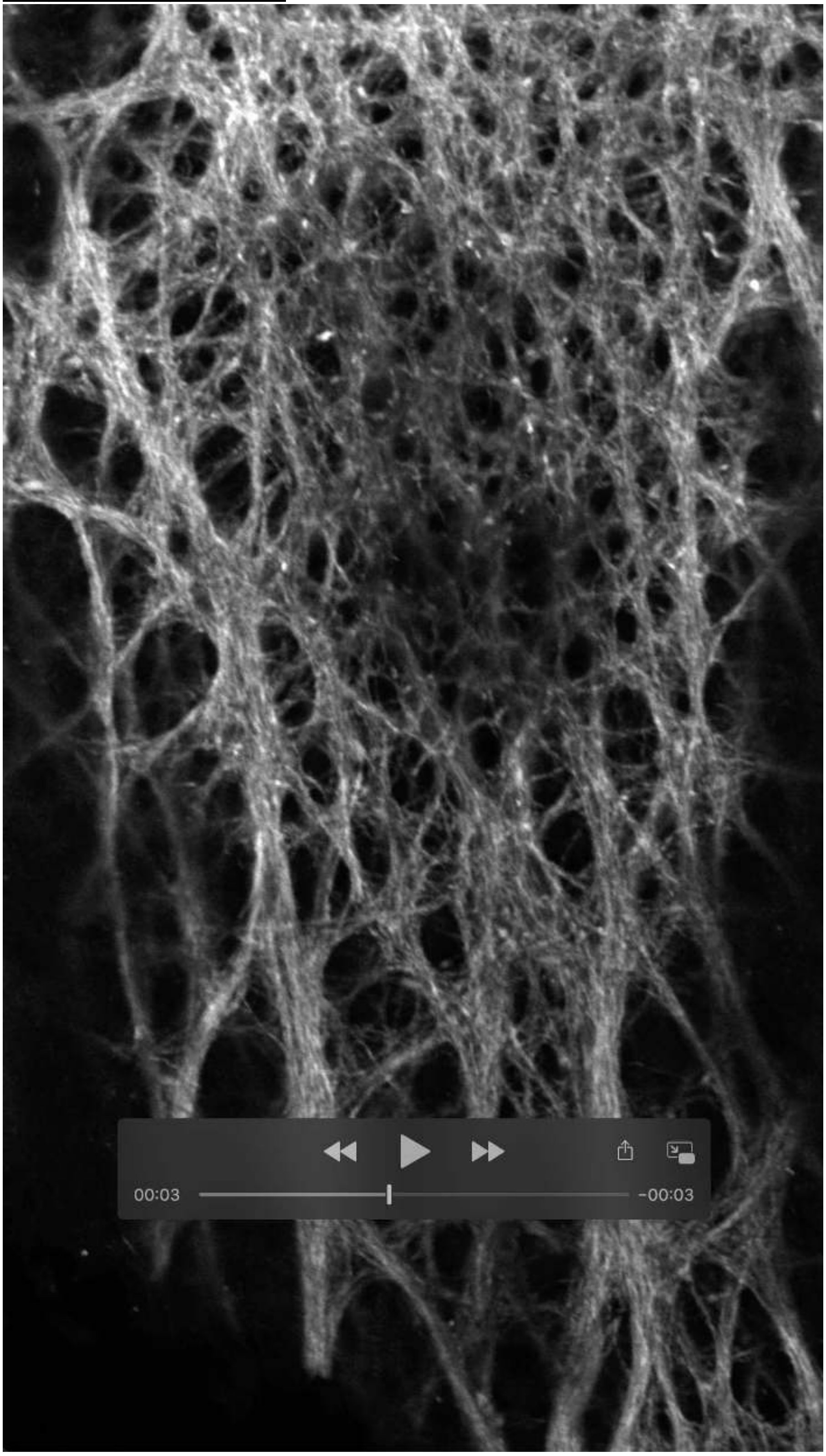

Supplementary Movie 2

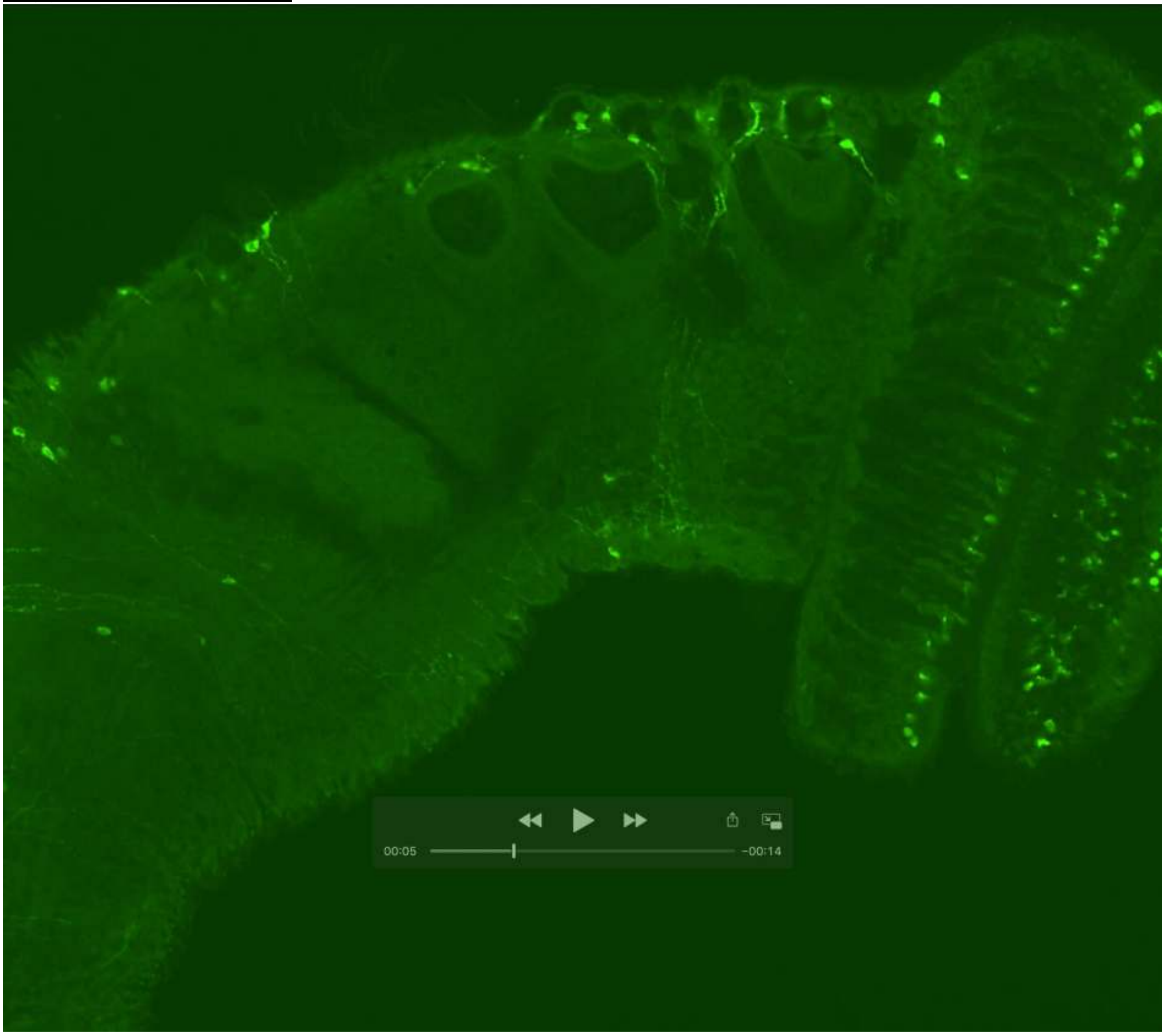
